## Supplementary Figures 1 to 7 for "An intrinsically disordered region mediates RNA-binding selectivity and pro-oncogenic activities of LARP6"

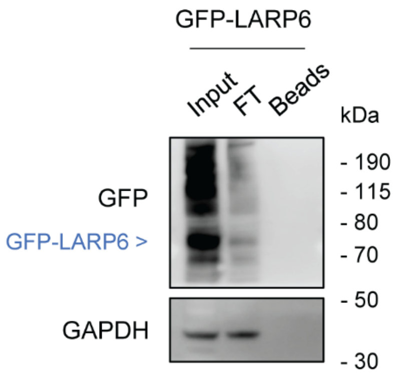


**Supplementary Figure 1: Confirmation of GFP-LARP6 immunoprecipitation for IPOOPS analysis.**

MDA-MB231 cells stably expressing GFP-LARP6 were subjected to UV-C crosslinking, followed by lysis and clearance of lysates with centrifugation. Aliquots were collected from the lysates before the IP (input), from the flowthrough (FT), and from the beads after the elution of GFP-LARP6 (beads) to validate the efficiency of the pulldown. The expected molecular weight of GFP-LARP6 is indicated in blue. The smear above the GFP-LARP6 band corresponds to the LARP6 portion crosslinked to RNA.

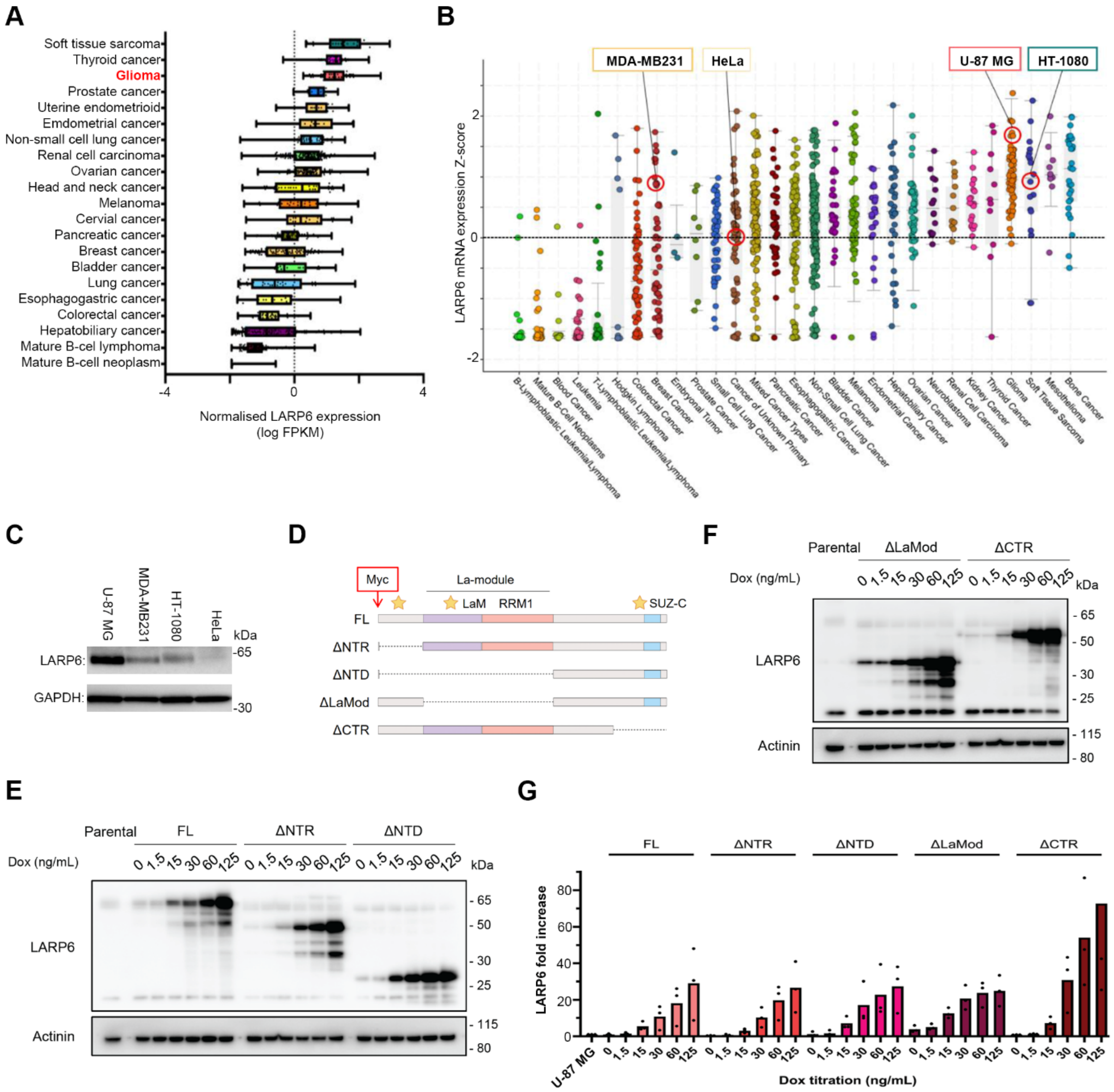


**Supplementary Figure 2: A doxycycline inducible U-87 MG glioblastoma cell-line model for characterisation of different LARP6 RNA binding regions in cells.**

**A)** LARP6 RNA expression levels across various cancer types from the TCGA database (Campbell et al., 2020) was extracted and exported from cBioPortal (cbioportal.org). Normalised LARP6 expression is displayed as log FPKM (fragment per kilobase of transcript per million).

**B)** LARP6 RNA expression values across various cancer cell lines. The expression data is from the Broad Institute Cancer Cell Line Encyclopaedia (Ghandi et al., 2019) and was exported from cBioPortal. Normalised LARP6 expression is displayed as z-scores relative to all samples in log RPKM (read per kilobase per million mapped reads). Cell lines are grouped according to their primary tumour site. The selected investigated cell lines are highlighted with red circles.

**C)** LARP6 protein expression levels across indicated cell lines. LARP6 protein expression was evaluated by Western blotting in the indicated cell lines of different cancer origins, chosen based on (B). Cancer origin: U-87 MG: glioblastoma; MDA-MB-231: triple negative breast adenocarcinoma; HT-1080: fibrosarcoma; HeLa: cervical cancer.

**D)** Schematic representation of myc-LARP6 inducible deletion mutants expressed in U-87 MG glioblastoma cells. All mutants express a myc-tag at the N-terminus.

**E)** Representative Western blot analysis of FL, ∆NTR, and ∆NTD myc-LARP6 deletion mutants induction of expression with different doses of doxycycline. Each myc-LARP6 variant was induced with 0–125 ng/mL doxycycline for 24 hours, before lysis and Western blot analysis with the indicated antibodies.

**F)** Representative Western blot analysis of ∆La-Module, and ∆CTR myc-LARP6 deletion mutants induction of expression with different doses of doxycycline. Each myc-LARP6 variant was induced with 0–125 ng/mL doxycycline for 24 hours, before lysis and Western blot analysis with the indicated antibodies.

**G)** Quantification of LARP6 expression levels from (E) and (F). LARP6 values were normalised to the endogenous LARP6 in parental U-87 MG cells.

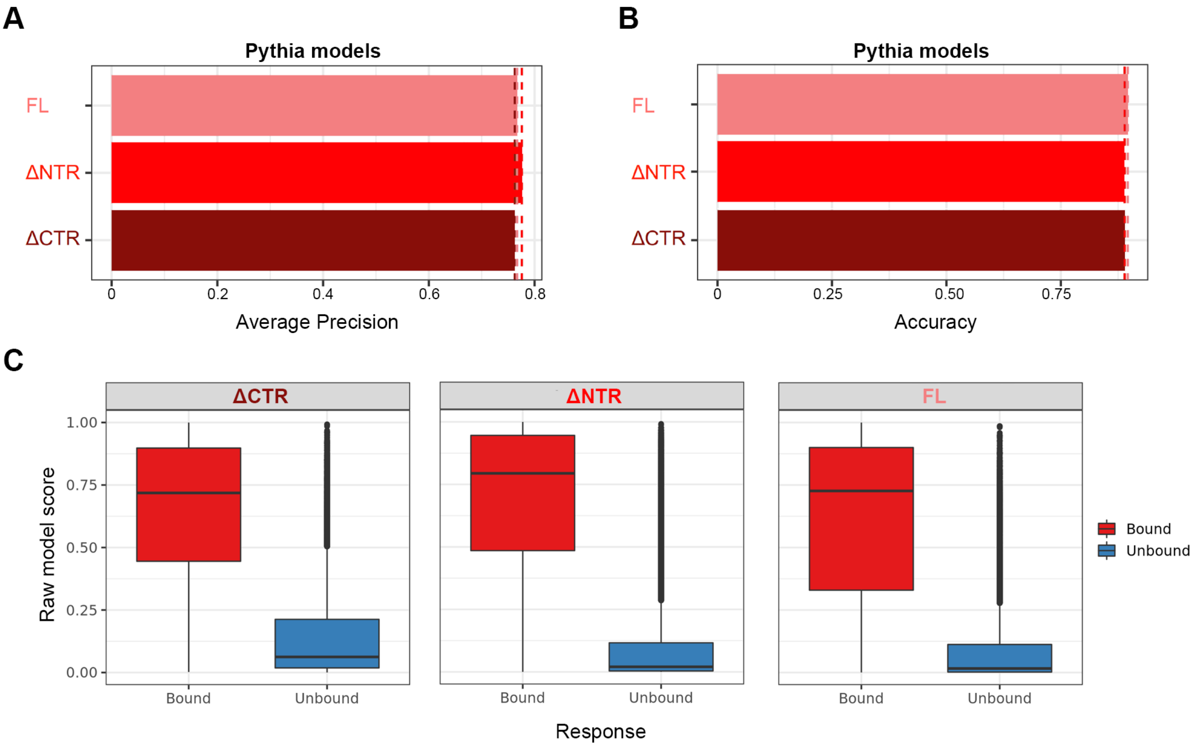


**Supplementary Figure 3: Benchmarking of Pythia for analysis of LARP6 wild-type and mutant RNA binding profiles.**

**A)** Average precision calculated for Pythia models trained on the FL, ΔNTR, and ΔCTR myc-LARP6 iCLIP peak datasets.

**B)** Average accuracy calculated for Pythia models trained on the FL, ΔNTR, and ΔCTR myc-LARP6 iCLIP peak datasets.

**C)** Raw model scores for predicting bound vs unbound peak fractions for Pythia models trained on the FL, ΔNTR, and ΔCTR myc-LARP6 iCLIP peak datasets.

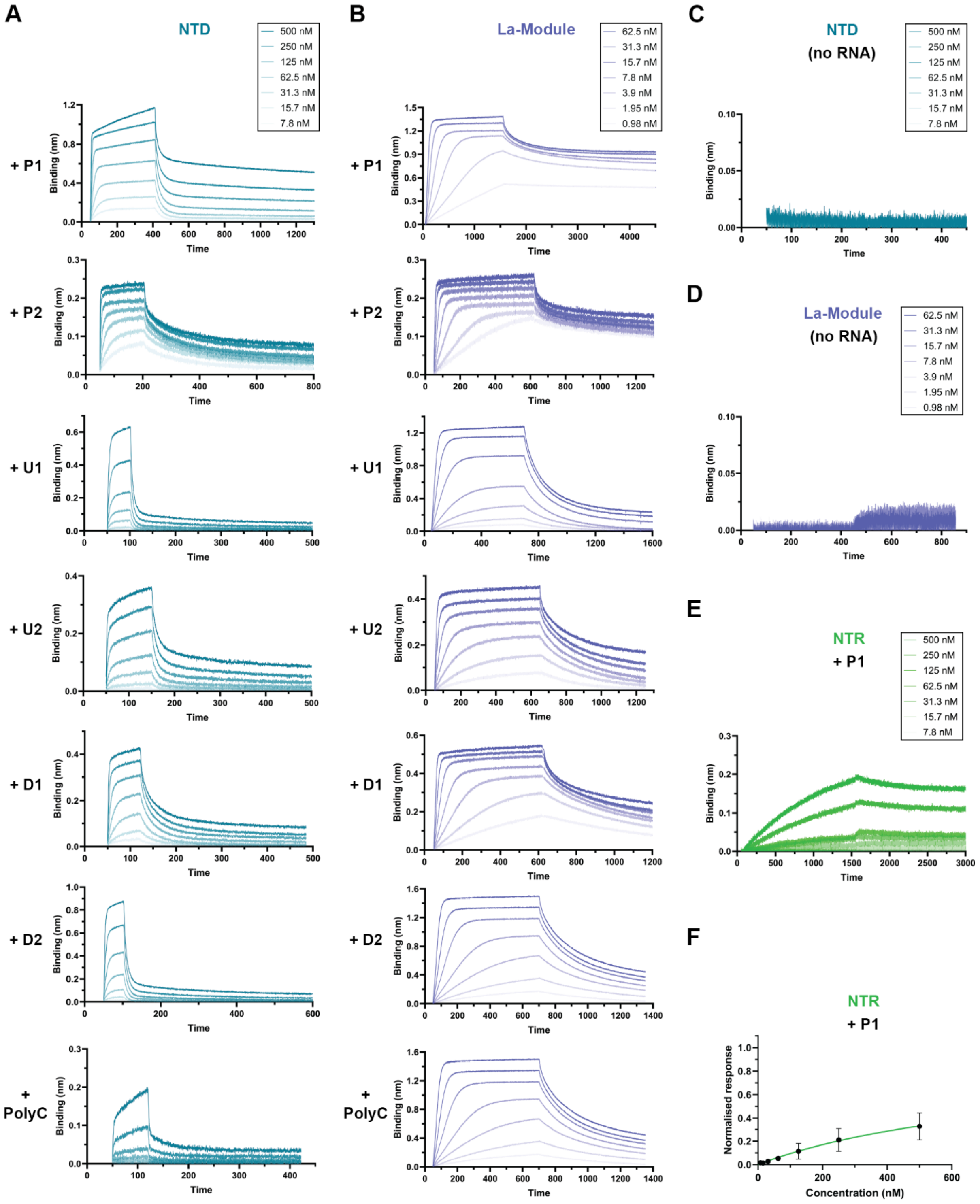


**Supplementary Figure 4: BLI analysis of the *in vitro* RNA binding activity of different LARP6 regions.**

**A)** Representative BLI association-dissociation profiles of the NTD segment of LARP6, against the indicated RNA oligonucleotides. The protein concentrations used for each plot are displayed within the top right box.

**B)** Representative BLI association-dissociation profiles of the La-module segment of LARP6, against the indicated RNA oligonucleotides. The protein concentrations used for each plot are displayed within the top right box.

**C)** BLI association-dissociation profile of the NTD segment of LARP6 in absence of RNA, as control for unspecific binding to empty sensors.

**D)** BLI association-dissociation profile of the La-Module segment of LARP6 in absence of RNA, as control for unspecific binding to empty sensors.

**E)** Representative association-dissociation profile of BLI experiments performed with the NTR segment of LARP6 and the P1 CTNNA1 3'UTR RNA oligonucleotide.

**F)** Steady state BLI binding analysis of the NTR segment of LARP6 and the P1 CTNNA1 3'UTR RNA oligonucleotide.

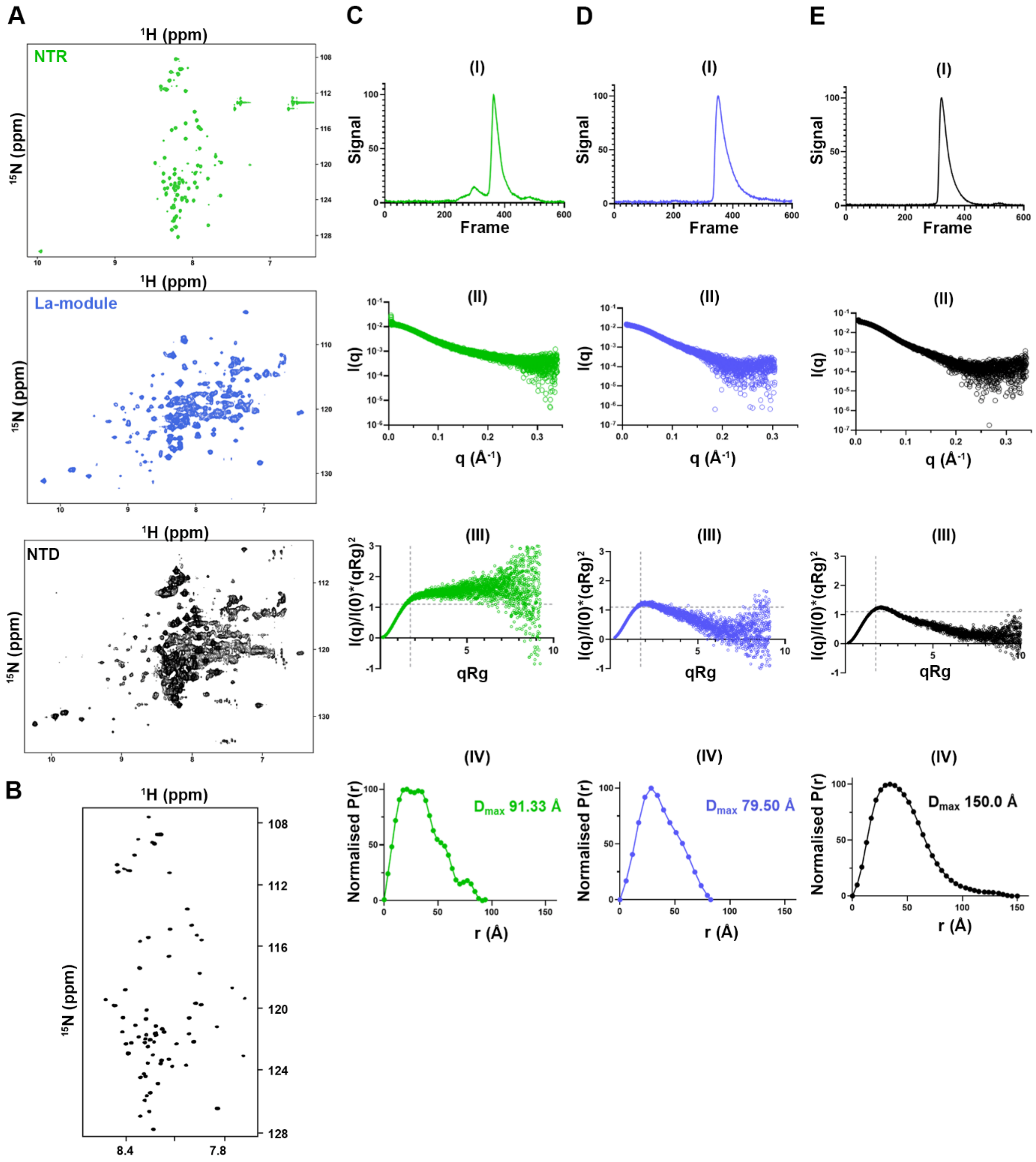


**Supplementary Figure 5: Biophysical characterisation of LARP6 NTR, La-module, and NTD segments in apo state.**

**A)** Full $^{1}H$-$^{15}N$ TROSY NMR spectra of LARP6 NTR, La-module, and NTD.

**B)** Spectrum of $^{1}H$-$^{15}N$ NOE experiment (saturated − 0.84 × control) for LARP6 NTR showing those amide groups experiencing fast internal motion.

**C)** SEC-SAXS characterisation of LARP6 NTD. The normalised signal of the integral ratio to the background for each frame of SEC-SAXS (I), the X-ray scattering curve obtained after buffer normalisation and averaging (II), the dimensionless Kratky plot (III) and the P(r) distance distribution plot (IV) are depicted.

**D)** SEC-SAXS characterisation of LARP6 La-module. The normalised signal of the integral ratio to the background for each frame of SEC-SAXS (I), the X-ray scattering curve obtained after buffer normalisation and averaging (II), the dimensionless Kratky plot (III), and the P(r) distance distribution plot (IV) are depicted.

**E)** SEC-SAXS characterisation of LARP6 NTR. The normalised signal of the integral ratio to the background for each frame of SEC-SAXS (I), the X-ray scattering curve obtained after buffer normalisation and averaging (II), the dimensionless Kratky plot (III), and the P(r) distance distribution plot (IV) are depicted.

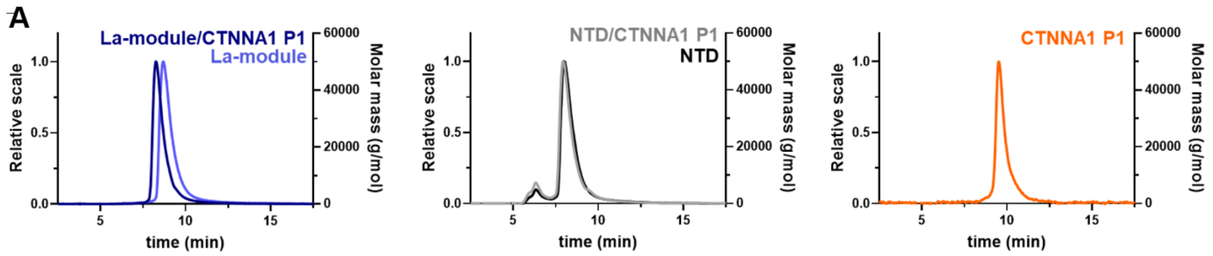


| Construct | Expected MW | Observed MW |
|---|---|---|
| La-module | 26.9 kDa | 36.4 ± 2.4 kDa |
| La-module/CTNNA1 P1 | 34.8 kDa | 46.3 ± 1.0 kDa |
| NTD | 33.9 kDa | 45.2 ± 0.5 kDa |
| NTD/CTNNA1 P1 | 41.8 kDa | 47.9 ± 0.3 kDa |

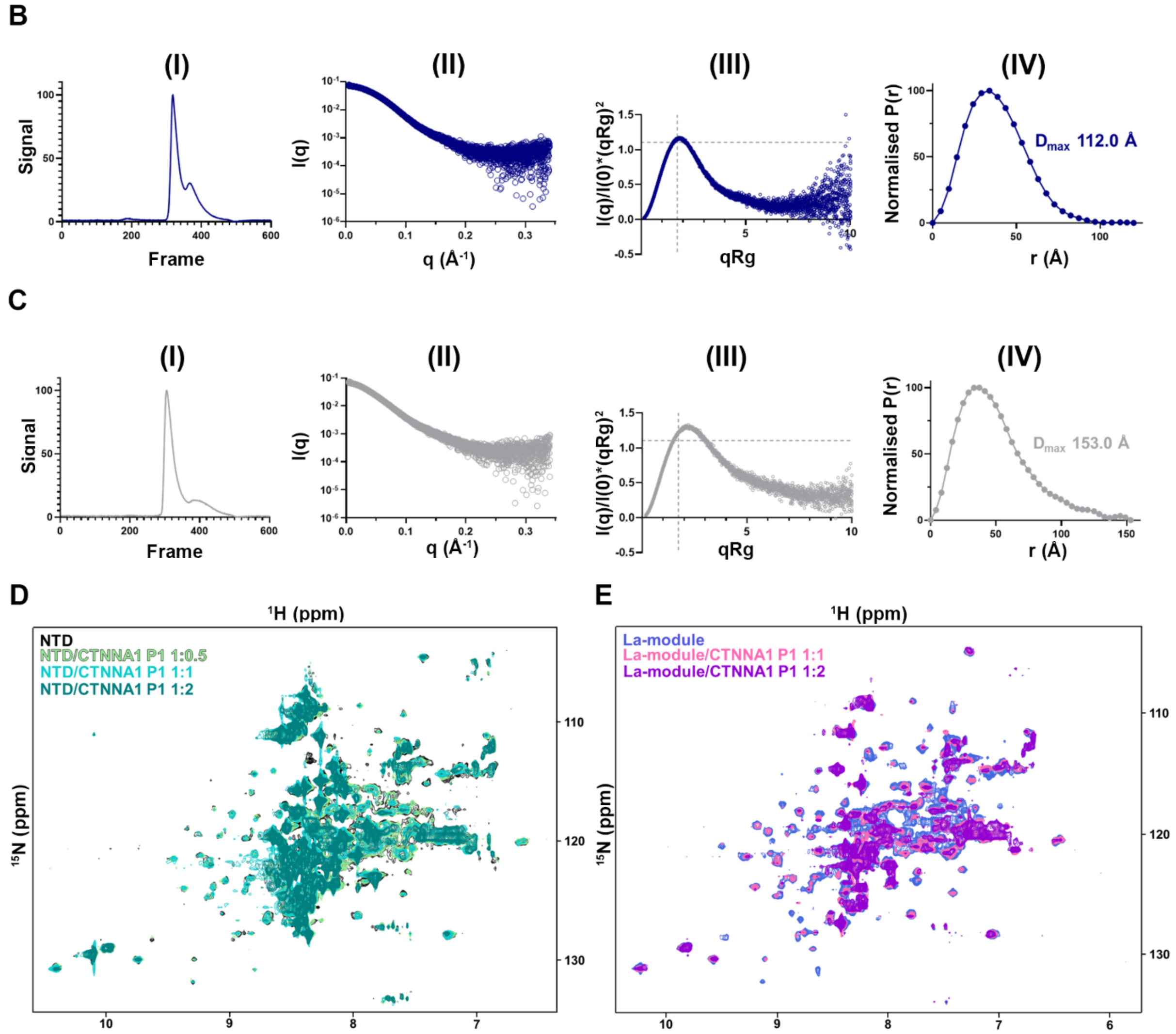


**Supplementary Figure 6: Biophysical characterisation of LARP6 La-module and NTD segments in RNA-bound state.**

**A)** SEC-MALS plots of La-module (left) and NTD (middle) in their apo and RNA bound states. The right panel shows the plot of the free CTNNA1 P1 RNA. The table reports the comparison between observed and expected molecular weights, based on theoretical masses calculation in Expasy ProtParam. The slightly higher masses obtained are consistent with the non-globular shape of the protein. The observed molecular weights of the complexes are compatible with a protein-RNA stoichiometry of 1:1.

**B)** SEC-SAXS characterisation of LARP6 La-module in complex with P1 RNA. The normalised signal of the integral ratio to the background for each frame of SEC-SAXS (I), the X-ray scattering curve obtained after buffer normalisation and averaging (II), the dimensionless Kratky plot (III), and the P(r) distance distribution plot (IV) are depicted.

**C)** SEC-SAXS characterisation of LARP6 NTD in complex with P1 RNA. The normalised signal of the integral ratio to the background for each frame of SEC-SAXS (I), the X-ray scattering curve obtained after buffer normalisation and averaging (II), the dimensionless Kratky plot (III), and the P(r) distance distribution plot (IV) from the analysis are depicted.

**D)** Overlay of the full $^{1}H$-$^{15}N$ TROSY NMR spectra of LARP6 NTD in the apo state and upon addition of the indicated molar ratios of P1 RNA.

**E)** Overlay of the full $^{1}H$-$^{15}N$ TROSY NMR spectra of LARP6 La-module in the apo state and upon addition of the indicated molar ratios of P1 RNA.

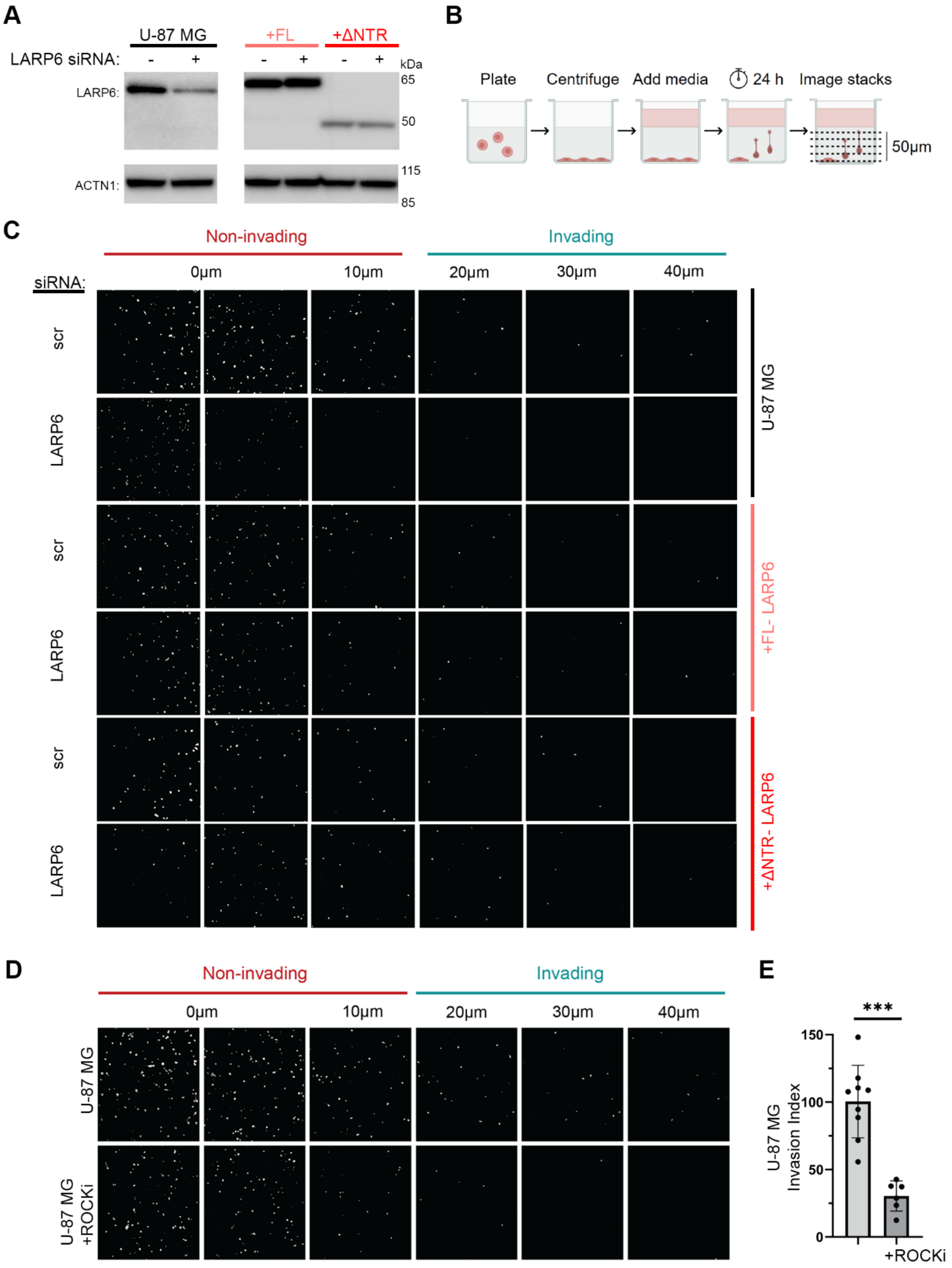


**Supplementary Figure 7: Knockdown-rescue analysis of cell invasion by LARP6.**

**A)** Representative Western blot analysis of endogenous LARP6 knockdown in parental U-87 MG cells, and rescue with the indicated myc-LARP6 variants.

**B)** Experimental workflow of 3D collagen invasion assay. Cells are embedded at the bottom of a collagen-I matrix mixture before allowing it to set into a gel. The cells are then induced to invade upwards into the collagen-I matrix for 24 hours by the addition of serum-containing media to the top of the gel, before fixation and analysis of the number of invaded cells by confocal microscopy.

**C)** Representative images of the invasion assay from Figure 5C. Nuclei are shown in white and were used at each z-plane to count the number of non-invaded vs. invaded cells.

**D)** Representative images of invasion assay in parental cells with or without 1μM AT13148 Rho kinase inhibitor (ROCKi). Nuclei are shown in white and were used at each z-plane to count the number of non-invaded vs. invaded cells.

**E)** Invasion index quantification from (D). The invasion index is normalised to the untreated control samples without ROCKi. Statistical test: Kolmogorov–Smirnov test (*** $p \leq 0.001$).
